# Naturally enhanced neutralizing breadth to SARS-CoV-2 after one year

**DOI:** 10.1101/2021.05.07.443175

**Authors:** Zijun Wang, Frauke Muecksch, Dennis Schaefer-Babajew, Shlomo Finkin, Charlotte Viant, Christian Gaebler, Hans-Heinrich Hoffmann, Christopher O. Barnes, Melissa Cipolla, Victor Ramos, Thiago Y. Oliveira, Alice Cho, Fabian Schmidt, Justin da Silva, Eva Bednarski, Lauren Aguado, Jim Yee, Mridushi Daga, Martina Turroja, Katrina G. Millard, Mila Jankovic, Anna Gazumyan, Zhen Zhao, Charles M. Rice, Paul D. Bieniasz, Marina Caskey, Theodora Hatziioannou, Michel C. Nussenzweig

**Affiliations:** Laboratory of Molecular Immunology, The Rockefeller University, New York, NY 10065, USA; Laboratory of Retrovirology, The Rockefeller University, New York, NY 10065, USA; Laboratory of Virology and Infectious Disease, The Rockefeller University, New York, NY, 10065, USA; Division of Biology and Biological Engineering, California Institute of Technology, Pasadena, CA, USA; Department of Pathology and Laboratory Medicine, Weill Cornell Medicine, New York, NY, 10065, USA; Howard Hughes Medical Institute

## Abstract

Over one year after its inception, the coronavirus disease-2019 (COVID-19) pandemic caused by severe acute respiratory syndrome coronavirus-2 (SARS-CoV-2) remains difficult to control despite the availability of several excellent vaccines. Progress in controlling the pandemic is slowed by the emergence of variants that appear to be more transmissible and more resistant to antibodies^1,2^. Here we report on a cohort of 63 COVID-19-convalescent individuals assessed at 1.3, 6.2 and 12 months after infection, 41% of whom also received mRNA vaccines^3,4^. In the absence of vaccination antibody reactivity to the receptor binding domain (RBD) of SARS-CoV-2, neutralizing activity and the number of RBD-specific memory B cells remain relatively stable from 6 to 12 months. Vaccination increases all components of the humoral response, and as expected, results in serum neutralizing activities against variants of concern that are comparable to or greater than neutralizing activity against the original Wuhan Hu-1 achieved by vaccination of naïve individuals^2,5-8^. The mechanism underlying these broad-based responses involves ongoing antibody somatic mutation, memory B cell clonal turnover, and development of monoclonal antibodies that are exceptionally resistant to SARS-CoV-2 RBD mutations, including those found in variants of concern^4,9^. In addition, B cell clones expressing broad and potent antibodies are selectively retained in the repertoire over time and expand dramatically after vaccination. The data suggest that immunity in convalescent individuals will be very long lasting and that convalescent individuals who receive available mRNA vaccines will produce antibodies and memory B cells that should be protective against circulating SARS-CoV-2 variants.

---

We initially characterized immune responses to SARS-CoV-2 in a cohort of convalescent individuals 1.3 and 6.2 months after infection^3,4^. Between February 8 and March 26, 2021, 63 participants between the ages of 26 and 73 years old (median 47 years) returned for a 12-month follow-up visit. Among those, 26 (41%) had received at least one dose of either the Moderna (mRNA-1273) or Pfizer-BioNTech (BNT162b2) vaccines, on average 40 days (range 2-82 days) before their study visit and 311 days (range 272-373 days) after onset of acute illness (Supplementary Table 1). Participants were was almost evenly split between sexes (43% female) and of the individuals that returned for a 12-month follow-up, only 10% had been hospitalized and the remainder had experienced relatively mild initial infections. Only 14% of the individuals reported persistent long-term symptoms after 12 months, reduced from 44% at the 6-month time point^4^. Symptom persistence was not associated with the duration and severity of acute disease or with vaccination status (Extended Data Fig. 1 a-c). All participants tested negative for active infection at the 12-month time point as measured by a saliva-based PCR assay ^4^. The demographics and clinical characteristics of the participants are shown in Supplementary Tables 1 and 2.

## Plasma SARS-CoV-2 Antibody Reactivity

Antibody reactivity in plasma to the RBD and nucleoprotein (N) were measured by enzyme-linked immunosorbent assay (ELISA)^3^. We limited our analysis to RBD because plasma anti-RBD antibodies are strongly correlated with neutralizing activity ^3,10-12^. Convalescent participants who had not been vaccinated maintained most of their anti-RBD IgM (103%), IgG (82%), and IgA (72%) titers between 6 and 12 months (Fig. 1a and Extended Data Fig. 2a-k). Consistent with previous reports^5–8^, vaccination increased the anti-RBD plasma antibody levels, with IgG titers increasing by nearly 30-fold compared to unvaccinated individuals (Fig. 1a right). The 2 individuals who did not show an increase had been vaccinated only 2 days before sample collection. In contrast to anti-RBD antibody titers that were relatively stable, anti-N antibody titers decreased significantly between 6 and 12 months in this assay irrespective of vaccination (Fig. 1b and Extended Data Fig. 2l-n).

**Fig. 1:**
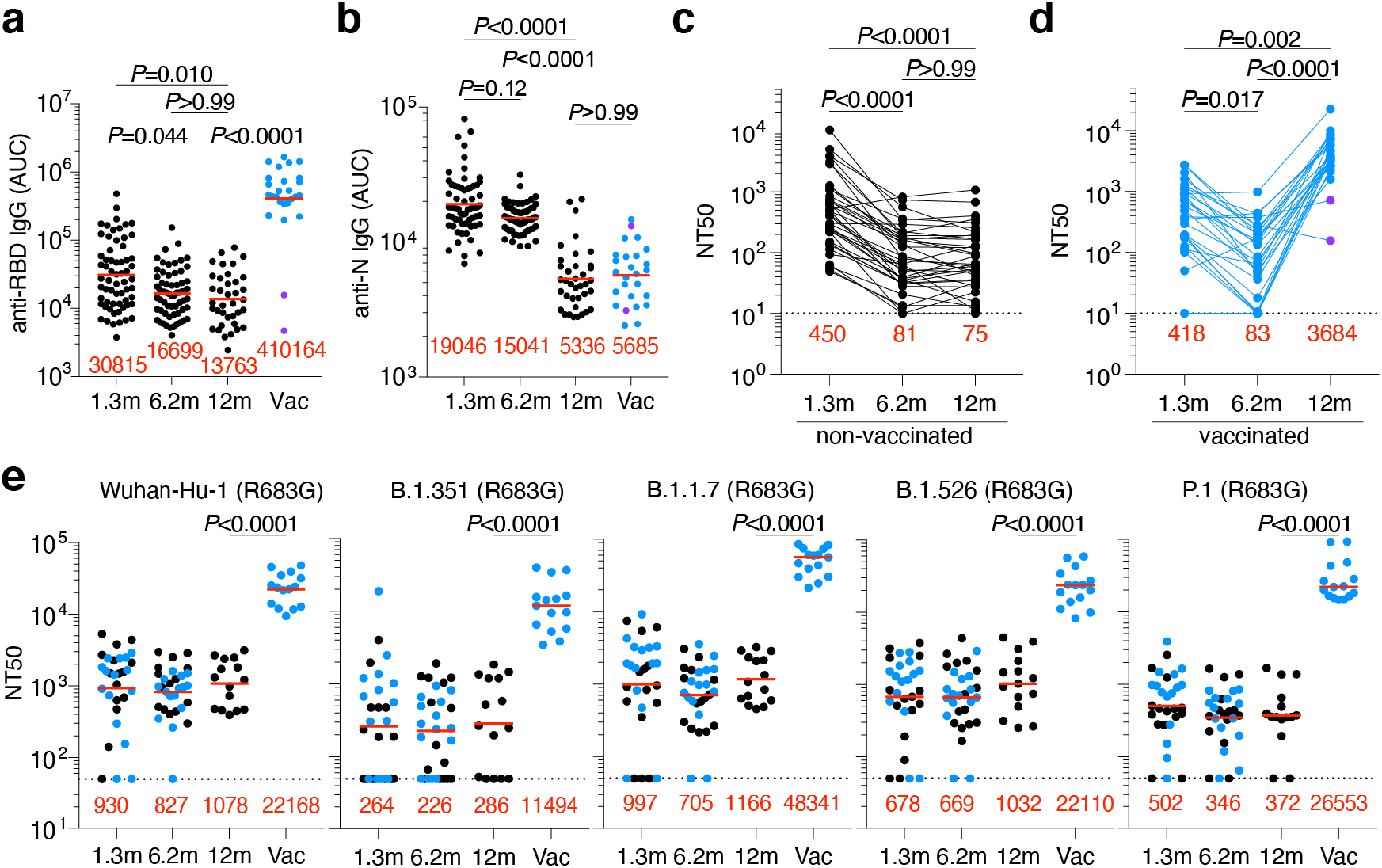
Plasma ELISAs and neutralizing activity. **a-b,** Plasma IgG antibody binding to SARS-CoV-2 RBD (**a**) and N protein (**b**), and plasma neutralizing activity (**c**–**d**) 12 months after infection (n=63). **a** and **b**, Area under the curve (AUC) over time in non-vaccinated and vaccinated individuals, as indicated. Numbers in red indicate geometric mean AUC at the indicated timepoint. n=63, 37 convalescent and 26 convalescent vaccinated individuals. Statistical significance in **a** and **b** was determined using two-sided Kruskal-Wallis test with subsequent Dunn’s multiple comparisons. **c, d,** NT50 over time in non-vaccinated (**c**) and vaccinated individuals (**d**). Lines connect longitudinal samples from the same individual. Statistical significance in **c-d** was determined using two-sided Friedman test with subsequent Dunn’s multiple comparisons. Two individuals who received their first dose of vaccine 24-48 hours before sample collection are depicted in purple. **e**, Plasma neutralizing activity against indicated SARS-CoV-2 variants of concern (n=30, 15 convalescent and 15 convalescent vaccinated individuals). The B.1.526 variant used here does contain the E484K substitution. Please refer to Methods for a detailed list of all substitutions/deletions/insertions of spike variants used here. Convalescent and convalescent vaccinated individuals in **a-e** are shown in black and blue, respectively. Statistical significance was determined using two-tailed Mann-Whitney test. Red numbers in c-e indicate the geometric mean NT50 at the indicated timepoint. All experiments were performed at least in duplicate.

Plasma neutralizing activity in 63 participants was measured using an HIV-1 pseudotyped with the SARS-CoV-2 spike protein ^3,4,13^ (Fig. 1c-d and Extended Data Fig. 2o). Twelve-months after infection, the geometric mean half-maximal neutralizing titer (NT_50_) for the 37 individuals that had not been vaccinated was 75, which was not significantly different from the same individuals at 6.2 months (Fig. 1c). In contrast, the vaccinated individuals showed a geometric mean NT_50_ of 3,684, which was nearly 50-fold greater than unvaccinated individuals and slightly better compared to the 30-fold increase in anti-RBD IgG antibodies (Fig. 1a, c, and d). Neutralizing activity was directly correlated with IgG anti-RBD (Extended Data Fig.2p) but not with anti-N titers (Extended Data Fig. 2r). We conclude that neutralizing titers remain relatively unchanged between 6 to 12 months after SARS-CoV-2 infection, and that vaccination further boosts this activity by nearly 50-fold.

To determine the neutralizing activity against circulating variants of concern/interest, we performed neutralization assays on HIV-1 virus pseudotyped with the S protein of the following SARS-CoV-2 variants of concern/interest: B.1.1.7, B.1.351, B.1.526 and P.1 ^1,14,15^. Twelve-months after infection neutralizing activity against the variants was generally lower than against wild-type SARS-CoV-2 virus in the same assay with the greatest loss of activity against B.1.351 (Fig. 1e). After vaccination the geometric mean NT_50_ rose to 11,493, 48,341, 22,109 and 26,553 against B.1.351, B.1.1.7, B.1.526 and P.1, respectively. These titers are an order of magnitude higher than the neutralizing titers we and others have reported against the wild-type SARS-CoV-2 at the peak of the initial response in infected individuals and in naïve individuals receiving both doses of mRNA vaccines (Fig. 1d and ^2–8^). Similar results were also obtained using authentic SARS-CoV-2 WA1/2020 and B.1.351 (Extended Data Fig. 2s).

## Memory B cells

The memory B cell compartment serves as an immune reservoir that contains a diverse collection of antibodies^16,17^. Although antibodies to the N-terminal domain and other parts of S can also be neutralizing, we limited our analysis to memory B cells that produce anti-RBD antibodies because they are the most numerous and potent ^18,19^. To enumerate RBD-specific memory B cells, we performed flow cytometry using a biotin-labeled RBD^3^ (Fig. 2a and Extended Data Fig. 3a and b). In the absence of vaccination, the number of RBD specific memory B cells present at 12 months was only 1.35-fold lower than the earlier timepoint (p= 0.027, Fig. 2a). In contrast and consistent with previous reports^5,8,20^, convalescent individuals that received mRNA vaccines showed an average 8.6-fold increase in the number of circulating RBD specific memory B cells (Fig. 2a). B cells expressing antibodies that bound to both wild-type and K417N/E484K/N501Y mutant RBDs were also enumerated by flow cytometry (Extended Data Fig. 3c). The number of variant RBD cross-reactive B cells was directly proportional to but 1.6 to 3.2-fold lower than wild-type RBD binding B cells (Fig. 2a).

**Fig. 2:**
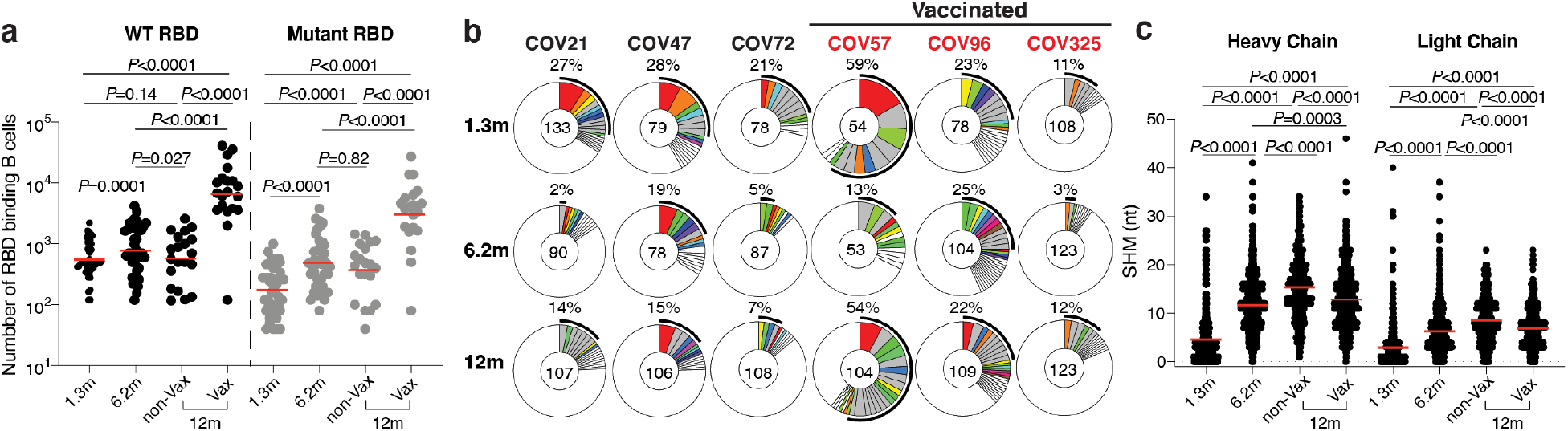
Anti-SARS-CoV-2 RBD B cell memory. **a,** graph summarizes number of antigen binding memory B cells per 2 million B cells (Extended data Fig 5b and c) obtained at 1.3, 6.2 and 12 months from 40 randomly selected individuals (vaccinees n=20, and non-vaccinees, n=20). Each dot is one individual. Red horizontal bars indicate geometric mean values. Statistical significance was determined using two-sided Kruskal-Wallis test with subsequent Dunn’s multiple comparisons. **b,** Pie charts show the distribution of antibody sequences from 6 individuals after 1.3 ^3^ (upper panel) or 6.2 ^4^ (middle panel) or 12 months (lower panel). The number in the inner circle indicates the number of sequences analyzed for the individual denoted above the circle. Pie slice size is proportional to the number of clonally related sequences. The black outline indicates the frequency of clonally expanded sequences detected in each participant. Colored slices indicate persisting clones (same IGV and IGJ genes, with highly similar CDR3s) found at both timepoints in the same participant. Grey slices indicate clones unique to the timepoint. White indicates sequences isolated once, and white slices indicate singlets found at both timepoints. **c.** Number of somatic nucleotide mutations in the IGVH and IGVL in antibodies (also Supplementary table 3) obtained after 1.3 or 6.2 or 12 months (1.3m: n=889; 6.2m: n=975; 12m: n=1105, (non-vax: n=417; vax: n=688)). Red horizontal bars indicate mean values. Statistical significance was determined using two-sided Kruskal-Wallis test with subsequent Dunn’s multiple comparisons.

The memory B cell compartment accumulates mutations and undergoes clonal evolution over the initial 6 months after infection^4,9,21,22^. To determine whether the memory compartment continues to evolve between 6 and 12 months, we obtained 1105 paired antibody heavy and light chain sequences from 10 individuals that were also assayed at the earlier time points, 6 of which were vaccinated (Fig. 2b, Extended Data Fig 3d, Supplementary Table 3). There were few significant differences among the expressed IGHV and IGLV genes between vaccinated and un-vaccinated groups, or between the 1.3-, 6-month and 1 year time points (Extended Data Fig 4a-c)^3,4^. *IGHV3-30* and *IGHV3-53* remained over-represented irrespective of vaccination ^10,18^(Extended Data Fig 4a).

All individuals assayed at 12 months showed expanded clones of RBD-binding memory cells that expressed closely related IGHV and IGLV genes (Fig. 2b, Extended Data Fig 3d and e). The relative fraction of cells belonging to these clones varied from 7-54% of the repertoire with no significant difference between vaccinated and non-vaccinated groups. The overall clonal composition differed between 6 and 12 months in all individuals suggesting ongoing clonal evolution (Fig. 2b and Extended Data Fig 3d). Among the 89 clones found after 12 months, 61% were not previously detected and 39% were present at one of the earlier time points (Fig. 2b and Extended Data Fig 3d). In vaccinated individuals the increase in size of the memory compartment was paralleled by an increase in the absolute number of B cells representing all persistent clones (Extended Data Fig 5a). Thus, RBD specific memory B cell clones were re-expanded upon vaccination in all 6 convalescent individuals examined (Fig. 2b and Extended Data Fig 3d and Extended Data Fig 5a).

Somatic hypermutation of antibody genes continued between 6 and 12 months after infection (Fig. 2c). Slightly higher levels of mutation were found in individuals who had not been vaccinated compared to vaccinated individuals possibly due to recruitment of newly-formed memory cells into the expanded memory compartment of the vaccinated individuals (Fig. 2c, Extended Data Fig 5b). There was no significant difference in mutation between conserved and newly arising clones at the 12-month time point in vaccinated individuals (Extended Data Fig 5c). Moreover, phylogenetic analysis revealed that sequences found at 6 and 12 months were intermingled and similarly distant from their unmutated common ancestors (Extended Data Fig 6). We conclude that clonal re-expansion of memory cells in response to vaccination is not associated with additional accumulation of large numbers of somatic mutations as might be expected if the clones were re-entering and proliferating in germinal centers.

## Neutralizing Activity of Monoclonal Antibodies

To determine whether the antibodies obtained from memory B cells 12 months after infection bind to RBD we performed ELISAs (Fig. 3a). 174 antibodies were tested by ELISA including: 1. 53 that were randomly selected from those that appeared only once and only after 1 year; 2. 91 that appeared as expanded clones or singlets at more than one time point; 3. 30 representatives of newly arising expanded clones (Supplementary Tables 4 and 5). Among the 174 antibodies tested, 173 bound to RBD indicating that the flow cytometry method used to identify B cells expressing anti-RBD antibodies was efficient (Supplementary Tables 4 and 5). The geometric mean ELISA half-maximal concentration (EC_50_) of the antibodies obtained after 12 months was 2.6 ng/ml, which was significantly lower than at 6 months irrespective of vaccination and suggestive of an increase in affinity (Fig. 3a, Extended data Fig. 7 a and b and Supplementary Tables 4 and 5). Consistent with this observation there was an overall increase in the apparent avidity of plasma antibodies between 1.3 and 12 months ^3,4^(p<0.0001, Extended data Fig. 7c).

**Fig. 3:**
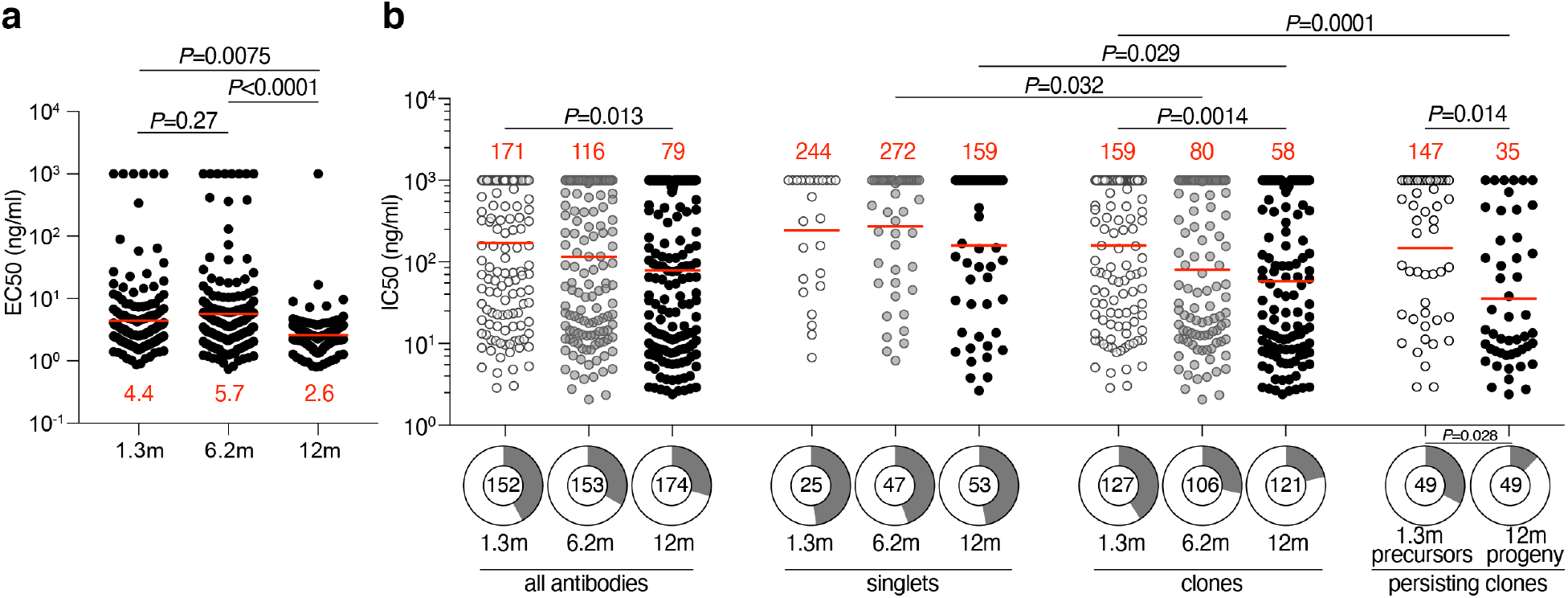
Anti-SARS-CoV-2 RBD monoclonal antibodies. **a,** Graph shows the ELISA binding EC_50_ (Y axis) for SARS-CoV-2 RBD by antibodies isolated at 1.3^3^ (n=152) 6.2^4^ (n=153) and 12 months (n=174) after infection. Statistical significance was determined using the two-sided Kruskal-Wallis test with subsequent Dunn’s multiple comparisons (1.3 vs 6.2 months, p=0.27; 1.3 vs 12 months, p=0.0075; 6.2 vs 12 months, p<0.0001). **b,** Graph shows anti-SARS-CoV-2 neutralizing activity of monoclonal antibodies measured by a SARS-CoV-2 pseudovirus neutralization assay^3,13^. Half-maximal inhibitory concentration (IC_50_) values for antibodies isolated at 1.3^3^ 6.2^4^ and 12 months after infection against wild-type SARS-CoV-2 (Wuhan-Hu-1 strain^41^) are shown. Each dot represents one antibody. Pie charts illustrate the fraction of non-neutralizing (IC_50_ > 1000 ng/ml) antibodies (grey slices), inner circle shows the number of antibodies tested per group. Horizontal bars and red numbers indicate geometric mean values. Statistical significance was determined through the two-sided Kruskal Wallis test with subsequent Dunn’s multiple comparisons.

All 174 RBD binding antibodies obtained from the 12-month time point were tested for neutralizing activity in a SARS-CoV-2 pseudotype neutralization assay. When compared to the earlier time points from the same individuals, the geometric mean half maximal inhibitory concentration (IC_50_) improved from 171 ng/mL (1.3 months) to 116 ng/mL (6 months) to 79 ng/mL (12 months), with no significant difference between vaccinated and non-vaccinated individuals (Fig. 3b and Extended data Fig. 7d, Supplementary Table 4). The increased potency was especially evident in the antibodies expressed by expanded clones of B cells that were conserved for the entire observation period irrespective of vaccination (p=0.014, Fig. 3b right, Extended Data Fig 7e-h and Supplementary Table 5). The overall increase in neutralizing activity among conserved clones was due to accumulation of clones expressing antibodies with potent neutralizing activity and simultaneous loss of clones expressing antibodies with no measurable activity (p= 0.028, Fig 3b right pie charts). Consistent with this observation, antibodies obtained from clonally expanded B cells after 12 months were more potent than antibodies obtained from unique B cells at the same time point (p= 0.029, Fig 3b).

## Epitopes and Breadth of Neutralization

To determine whether the loss of non-neutralizing antibodies over time was due to preferential loss of antibodies targeting specific epitopes, we performed BLI experiments in which a preformed antibody-RBD complex was exposed to a second monoclonal targeting one of 3 classes of structurally defined epitopes^3,23^ (see schematic in Fig. 4a). We assayed 60 randomly selected antibodies with comparable neutralizing activity from the 1.3- and 12-month time points. The 60 antibodies were evenly distributed between the 2 time points and between neutralizers and non-neutralizers (Fig. 4). Antibody affinities for RBD were similar among neutralizers and non-neutralizers obtained at the same time point (Fig. 4b, Extended Data Fig. 8). When the two sets of un-related antibodies obtained from 1.3 and 12 months were compared, they showed significantly increased affinity over time irrespective of their neutralizing activity (Fig. 4b, Extended Data Fig. 8). In competition experiments, all but 2 of the 30 non-neutralizing antibodies failed to inhibit binding of the class 1 (C105), 2 (C121 and C144) or 3 (C135) antibodies tested and therefore must bind to epitopes that do not overlap with the epitopes of these classes of antibodies (Fig. 4c, and Extended Data Fig. 9). In contrast, all but 2 of the 30 neutralizers blocked class 1, or 2 antibodies whose target epitopes are structural components of the RBD that interact with its cellular receptor, the angiotensin-converting enzyme 2^23,24^ (ACE2) (Fig. 4c and Extended Data Fig. 9). In addition, whereas 9 of the 15 neutralizing antibodies obtained after 1.3 months blocked both class 1 and 2 antibodies, only 1 of the 15 obtained after 12 months did so. In contrast to the earlier time point, 13 of 15 neutralizing antibodies obtained after 12 months only interfered with C121, a class 2 antibody^3,23^ (Fig. 4c and Extended Data Fig. 9). We conclude that neutralizing antibodies are retained and non-neutralizing antibodies targeting RBD surfaces that do not interact with ACE2 are removed from the repertoire over time.

**Fig. 4:**
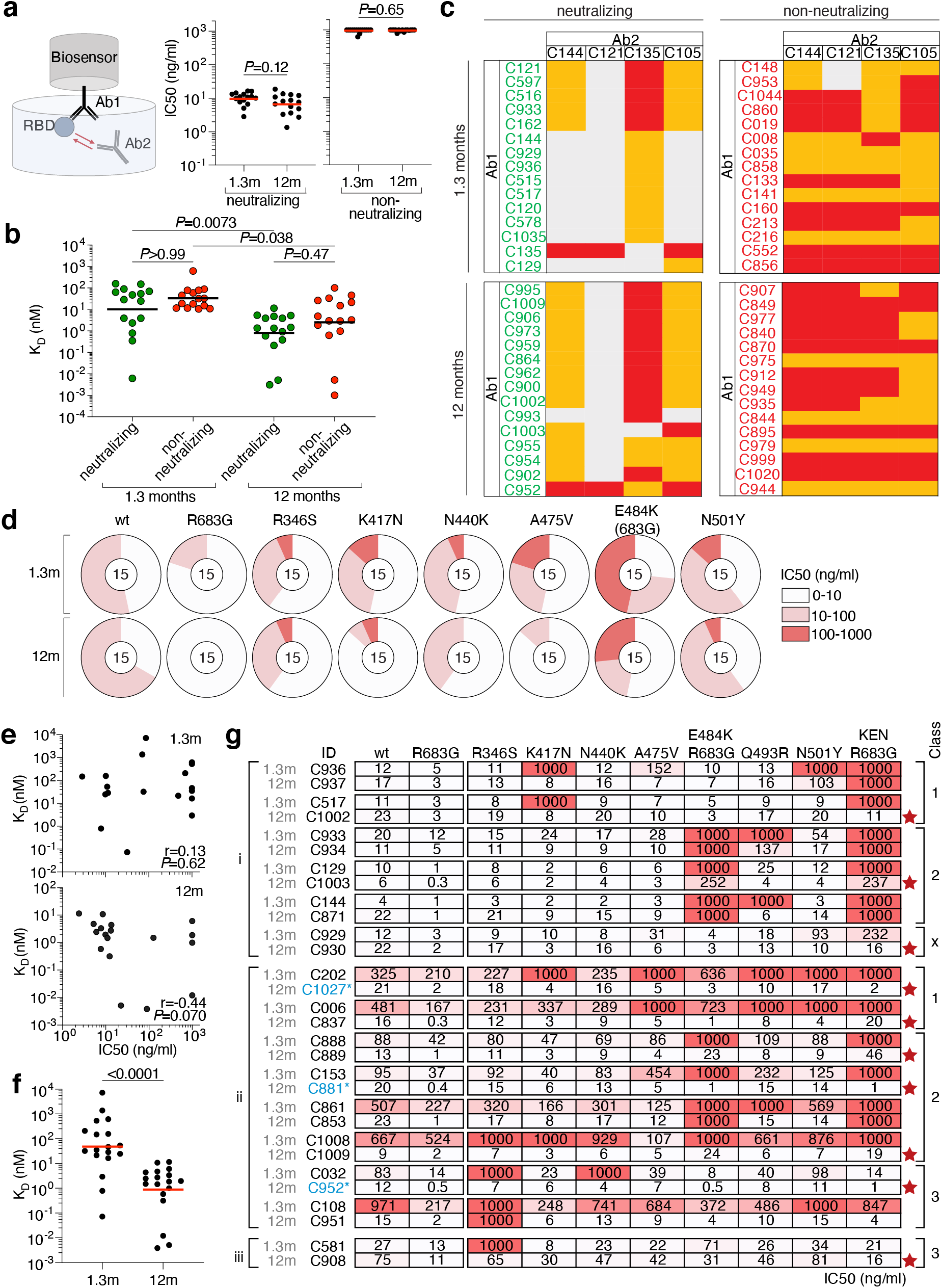
Epitope targeting and evolution of anti-SARS-CoV-2 RBD antibodies. **a,** Schematic representation of the BLI experiment (left) and IC_50_ values for randomly selected neutralizing (middle) and non-neutralizing (right) antibodies isolated at 1.3- and 12-months post-infection (each presented group shows n=15 antibodies, resulting in a total of n=60 antibodies). Red horizontal bars indicate geometric mean. Statistical significance was determined using two-sided Mann-Whitney test. **b**, K_D_ values of the n=30 neutralizing (green) and n=30 non-neutralizing (red) antibodies shown in a. Horizontal bars indicate geometric mean values. Statistical significance was determined using two-sided Kruskal Wallis test with subsequent Dunn’s multiple comparisons. BLI traces can be found in Extended data Fig. 8. **c,** Heat-map of relative inhibition of Ab2 binding to the preformed Ab1-RBD complexes (grey=no binding, orange=intermediate binding, red=high binding). Values are normalized through the subtraction of the autologous antibody control. BLI traces can be found in Extended Data Fig 9. **d**, Neutralization of the indicated mutants for antibodies shown in panel **a-c**. Pie charts illustrate the fraction of antibodies that are poorly/non-neutralizing (IC_50_ 100-1000 ng/ml, red), intermediate neutralizing (IC_50_ 10-100 ng/ml, pink) and potently neutralizing (IC_50_ 0-10 ng/ml, white) for each mutant. Number in inner circle shows number of antibodies tested. **e,** Graphs show affinities (Y axis) plotted against neutralization activity (X axis) for 18 clonal antibody pairs isolated 1.3 (top) and 12 months (bottom) (after infection (n=36 antibodies). Statistical significance was determined using Spearman correlation test. **f,** BLI affinity measurements for same n=36 paired 1.3- and 12-month antibodies as in **e**. Statistical significance was determined using two-tailed Wilcoxon test. **g,** IC_50_ values for n=30 paired neutralizing antibodies isolated at indicated timepoints against indicated mutant SARS-CoV-2 pseudoviruses. Antibodies are divided into groups i-iii, based on neutralizing activity: (i) potent clonal pairs that do not improve over time, (ii) clonal pairs that show increased activity over time, and (iii) and clonal pairs showing decreased neutralization activity after 12 months. Antibody class assignment based on initial (1.3m) sensitivity to mutation is indicated on the right. Red stars indicate antibodies that neutralize all RBD mutants tested. Color gradient indicates IC_50_ values ranging from 0 (white) to 1000 ng/ml (red).

To determine whether there was an increase in neutralization breadth over time, the neutralizing activity of the 60 antibodies was assayed against a panel of RBD mutants covering residues associated with circulating variants of concern: R346S, K417N, N440K, A475V, E484K and N501Y (Fig. 4d and Supplementary Table 6). Increased activity was evident against K417N, N440K, A475V, E484K and N501Y (Fig. 4d and Supplementary Table 6). We conclude that evolution of the antibody repertoire results in acquisition of neutralization breadth over time.

The increase in breadth and overall potency of memory B cell antibodies could be due to shifts in the repertoire, clonal evolution, or both. To determine whether changes in specific clones are associated with increases in affinity and breadth, we measured the relative affinity and neutralizing breadth of matched pairs of antibodies expressed by expanded clones of B cells that were maintained in the repertoire over the entire observation period^3,4^. SARS-CoV-2 neutralizing activity of the antibodies present at 1.3 or 12 months was not significantly correlated with affinity at either time point when each time point is considered independently (Fig. 4e).

However, there was a significant increase in overall affinity over time including in the 4 pairs of antibodies with no measurable neutralizing activity (Fig. 4f and Supplementary Table 7). Neutralizing breadth was assayed for 15 randomly selected pairs of antibodies targeting epitopes assigned to the 3 dominant classes of neutralizing antibodies^3,23,25,26^. Seven of the selected antibodies showed equivalent or decreased activity against wild-type SARS-CoV-2 after 12 months (Fig. 4g and Supplementary Table 8). However, neutralizing breadth increased between 1.3 and 12-months for all 15 pairs, even when neutralizing activity against the wild-type was unchanged or decreased (Fig. 4g and Supplementary Table 8). Only 1 of the 15 antibodies obtained after 1.3 months neutralized all the mutants tested (Fig. 4g). In contrast, 10 of the 15 antibodies obtained from the same clones after 12 months neutralized all variants tested with IC_50_s as low as 1 ng/ml against the triple mutant K417N/E484K/N501Y found in B.1.351 (Fig. 4g and Supplementary Table 8). Similar results were obtained with authentic WA1/2020 and B.1.351 (Extended Data Fig 7i). In conclusion, continued clonal evolution of anti-SARS-CoV-2 antibodies over 12 months favors increasing potency and breadth resulting in monoclonal antibodies with exceptional activity against a broad group of variants.

## Discussion

During immune responses activated B cells interact with cognate T cells and begin dividing before selection into the plasma cell, memory or germinal center B cell compartments based in part on their affinity for antigen^17,27-31^. Whereas B cells expressing high affinity antibodies are favored to enter the long-lived plasma cell compartment, the memory compartment is more diverse and can develop directly from activated B cells or from a germinal center^17,27-31^. Memory cells emanating from a germinal center carry more mutations than those that develop directly from activated B cells because they undergo additional cycles of division^32^.

Consistent with the longevity of bone marrow plasma cells, infection with SARS-CoV-2 leads to persistent serum anti-RBD antibodies, and corresponding neutralizing responses. Nearly 93% of the plasma neutralizing activity is retained between 6- and 12-months^33,34^. Vaccination boosts the neutralizing response by 1.5 orders of magnitude by inducing additional plasma cell differentiation from the memory B cell compartment ^5,7,35^. Recruitment of evolved memory B cells producing antibodies with broad and potent neutralizing activity into the plasma cell compartment is likely to account for the exceptional serologic activity of vaccinated convalescents against variants of concern^20,35,36^.

Less is known about selection and maintenance of the memory B cell compartment. SARS-CoV-2 infection produces a memory compartment that continues to evolve over 12 months after infection with accumulation of somatic mutations, emergence of new clones, and increasing affinity all of which is consistent with long-term persistence of germinal centers. The increase in activity against SARS-CoV-2 mutants parallels the increase in affinity and is consistent with the finding that increasing the apparent affinity of anti-SARS-CoV-2 antibodies by dimerization or by creating bi-specific antibodies also increases resistance to RBD mutations^37-40^.

Continued antibody evolution in germinal centers requires antigen which can be retained in these structures over long periods of time^29^. In addition, SARS-CoV-2 protein and nucleic acid has been reported in the gut for at least 2 months after infection^4^. Irrespective of the source of antigen, antibody evolution favors epitopes overlapping with the ACE2 binding site on the RBD, possibly because these are epitopes that are preferentially exposed on trimeric spike protein or virus particles.

Vaccination after SARS-CoV-2 infection increases the number of RBD binding memory cells by over an order of magnitude by recruiting new B cell clones into memory and expanding persistent clones. The persistent clones expand without accumulating large numbers of additional mutations indicating that clonal expansion of human memory B cells does not require re-entry into germinal centers and occurs through the activated B cell compartment^17,27,31^.

The remarkable evolution of breadth after infection and the robust enhancement of serologic responses and B cell memory achieved with mRNA vaccination suggests that convalescent individuals who are vaccinated should enjoy high levels of protection against emerging variants without a need to modify existing vaccines. If memory responses evolve in a similar manner in naive individuals that receive vaccines, additional appropriately timed boosting with available vaccines should lead to protective immunity against circulating variants.

## Methods

### Study participants

Previously enrolled study participants were asked to return for a 12-month follow-up visit at the Rockefeller University Hospital in New York from February 8 to March 26, 2021. Eligible participants were adults with a history of participation in both prior study visits of our longitudinal cohort study of COVID-19 recovered individuals^3,4^. All participants had a confirmed history of SARS-CoV-2 infection, either diagnosed during the acute infection by RT-PCR or retrospectively confirmed by seroconversion. Exclusion criteria included presence of symptoms suggestive of active SARS-CoV-2 infection. Most study participants were residents of the Greater New York City tri-state region and were asked to return approximately 12 months after the time of onset of COVID-19 symptoms. Participants presented to the Rockefeller University Hospital for blood sample collection and were asked about potential symptom persistence since their 6.2-month study visit, laboratory-confirmed episodes of reinfection with SARS-CoV-2, and whether they had received any COVID-19 related treatment or SARS-CoV-2 vaccination in the interim. Study participants who had received COVID-19 vaccinations, were exclusively recipients of one of the two currently EUA-approved mRNA vaccines, Moderna (mRNA-1273) or Pfizer-BioNTech (BNT162b2), and individuals who received both doses did so according to current interval guidelines, namely 28 days (range 28-30 days) for Moderna and 21 days (range 21-23 days) for Pfizer-BioNtech. Detailed characteristics of the symptomology and severity of the acute infection, symptom kinetics, and the immediate convalescent phase (7 weeks post-symptom onset until 6.2month visit) have been reported previously ^4^. Participants that presented with persistent symptoms attributable to COVID-19 were identified on the basis of chronic shortness of breath or fatigue, deficit in athletic ability and/or three or more additional long-term symptoms such as persistent unexplained fevers, chest pain, new-onset cardiac sequalae, arthralgias, impairment of concentration/mental acuity, impairment of sense of smell/taste, neuropathy or cutaneous findings as previously described ^4^. Clinical data collection and management were carried out using the software iRIS by iMedRIS. All participants at Rockefeller University provided written informed consent before participation in the study and the study was conducted in accordance with Good Clinical Practice. For detailed participant characteristics see Supplementary Table 2. The study was performed in compliance with all relevant ethical regulations and the protocol (DRO-1006) for studies with human participants was approved by the Institutional Review Board of the Rockefeller University.

### SARS-CoV-2 molecular tests

Saliva was collected into guanidine thiocyanate buffer as described ^42^. RNA was extracted using either a column-based (Qiagen QIAmp DSP Viral RNA Mini Kit, Cat#61904) or a magnetic bead-based method as described ^43^. Reverse transcribed cDNA was amplified using primers and probes validated by the CDC or by Columbia University Personalized Medicine Genomics Laboratory respectively and approved by the FDA under the Emergency Use Authorization. Viral RNA was considered detected if Ct for two viral primers/probes were <40.

### Blood samples processing and storage

Peripheral Blood Mononuclear Cells (PBMCs) obtained from samples collected at Rockefeller University were purified as previously reported by gradient centrifugation and stored in liquid nitrogen in the presence of FCS and DMSO ^3,4^. Heparinized plasma and serum samples were aliquoted and stored at −20 °C or less. Prior to experiments, aliquots of plasma samples were heat-inactivated (56 °C for 1 hour) and then stored at 4 °C.

### ELISAs

ELISAs^44,45^ to evaluate antibodies binding to SARS-CoV-2 RBD and N were performed by coating of high-binding 96-half-well plates (Corning 3690) with 50 μl per well of a 1μg/ml protein solution in PBS overnight at 4 °C. Plates were washed 6 times with washing buffer (1× PBS with 0.05% Tween-20 (Sigma-Aldrich)) and incubated with 170 μl per well blocking buffer (1× PBS with 2% BSA and 0.05% Tween-20 (Sigma)) for 1 h at room temperature. Immediately after blocking, monoclonal antibodies or plasma samples were added in PBS and incubated for 1 h at room temperature. Plasma samples were assayed at a 1:66 starting dilution and 7 (IgA and IgM anti-RBD) or 11 (IgG anti-RBD) additional threefold serial dilutions. Monoclonal antibodies were tested at 10 μg/ml starting concentration and 10 additional fourfold serial dilutions. Plates were washed 6 times with washing buffer and then incubated with anti-human IgG, IgM or IgA secondary antibody conjugated to horseradish peroxidase (HRP) (Jackson Immuno Research 109-036-088 109-035-129 and Sigma A0295) in blocking buffer at a 1:5,000 dilution (IgM and IgG) or 1:3,000 dilution (IgA). Plates were developed by addition of the HRP substrate, TMB (ThermoFisher) for 10 min (plasma samples) or 4 minutes (monoclonal antibodies). The developing reaction was stopped by adding 50 μl 1 M H_2_SO_4_ and absorbance was measured at 450 nm with an ELISA microplate reader (FluoStar Omega 5.11, BMG Labtech) with Omega MARS software for analysis. For plasma samples, a positive control (plasma from participant COV72, diluted 66.6-fold and seven additional threefold serial dilutions in PBS) was added to every assay plate for validation. The average of its signal was used for normalization of all of the other values on the same plate with Excel software before calculating the area under the curve using Prism V9.1(GraphPad). For monoclonal antibodies, the EC50 was determined using four-parameter nonlinear regression (GraphPad Prism V9.1).

### Proteins

Mammalian expression vectors encoding the RBDs of SARS-CoV-2 (GenBank MN985325.1; S protein residues 319-539) or K417N, E484K, N501Y RBD mutants with an N-terminal human IL-2 or Mu phosphatase signal peptide were previously described ^46^. SARS-CoV-2 nucleocapsid protein (N) was purchased from Sino Biological (40588-V08B).

### SARS-CoV-2 pseudotyped reporter virus

A panel of plasmids expressing RBD-mutant SARS-CoV-2 spike proteins in the context of pSARS-CoV-2-S_Δ19_ has been described previously ^2,9,26^. Variant pseudoviruses resembling variants of concern B.1.1.7 (first isolated in the UK), B.1.351 (first isolated in South-Africa), B.1.526 (first isolated in New York City) and P.1 (first isolated in Brazil) were generated by introduction of substitutions using synthetic gene fragments (IDT) or overlap extension PCR mediated mutagenesis and Gibson assembly. Specifically, the variant-specific deletions and substitutions introduced were:

B.1.1.7: ΔH69/V70, ΔY144, N501Y, A470D, D614G, P681H, T761I, S982A, D118H
B.1.351: D80A, D215G, L242H, R246I, K417N, E484K, N501Y, D614G, A701V
B.1.526: L5F, T95I, D253G, E484K, D614G, A701V.
P.1: L18F, R20N, P26S, D138Y, R190S, K417T, E484K, N501Y, D614G, H655Y

The E484K and K417N/E484K/N501Y (KEN) substitution, as well as the deletions/substitutions corresponding to variants of concern were incorporated into a spike protein that also includes the R683G substitution, which disrupts the furin cleaveage site and increases particle infectivity. Neutralizing activity against mutant pseudoviruses were compared to a wildtype SARS-CoV-2 spike sequence (NC_045512), carrying R683G where appropriate.

SARS-CoV-2 pseudotyped particles were generated as previously described ^3,13^. Briefly, 293T cells were transfected with pNL4-3ΔEnv-nanoluc and pSARS-CoV-2-S_Δ19_, particles were harvested 48 hpt, filtered and stored at −80°C.

### Microneutralization assay with authentic SARS-CoV-2

Microneutralization assay of SARS-CoV-2 virus were performed as described previously^3^. The day prior to infection, Vero E6 cells were seeded at 1X10^4^ cells/well into 96-well plates. The diluted plasma and antibodies were mixed with SARS-CoV-2 WA1/2020 or the SA variant B.1.351 and incubated for 1 hour at 37°C. The antibody-virus-mix was then directly applied to Vero E6 cells and incubated for 22 hours at 37°C. Cells were subsequently fixed by adding an equal volume of 70% formaldehyde to the wells, followed by permeabilization with 1% Triton X-100 for 10 minutes. After washing, cells were incubated for 1 hour at 37°C with blocking solution of 5% goat serum in PBS (catalog no. 005-000-121; Jackson ImmunoResearch). A rabbit polyclonal anti-SARS-CoV-2 nucleocapsid antibody (catalog no. GTX135357; GeneTex) was added to the cells at 1:1,000 dilution in blocking solution and incubated at 4 °C overnight. Goat anti-rabbit AlexaFluor 594 (catalog no. A-11012; Life Technologies) was used as a secondary antibody at a dilution of 1:2000. Nuclei were stained with Hoechst 33342 (catalog no. 62249; Thermo Fisher Scientific) at 1μg/ml. Images were acquired with a fluorescence microscope and analyzed using ImageXpress Micro XLS (Molecular Devices). All experiments were performed in a biosafety level 3 laboratory.

### Pseudotyped virus neutralization assay

Fourfold serially diluted plasma from COVID-19-convalescent individuals or monoclonal antibodies were incubated with SARS-CoV-2 pseudotyped virus for 1 h at 37 °C. The mixture was subsequently incubated with 293T_Ace2_ cells^3^ (for comparisons of plasma or monoclonal antibodies from convalescent individuals) or HT1080Ace2 cl14 cells^13^ (for analyses involving mutant/variant pseudovirus panels), as indicated, for 48h after which cells were washed with PBS and lysed with Luciferase Cell Culture Lysis 5× reagent (Promega). Nanoluc Luciferase activity in lysates was measured using the Nano-Glo Luciferase Assay System (Promega) with the Glomax Navigator (Promega). The obtained relative luminescence units were normalized to those derived from cells infected with SARS-CoV-2 pseudotyped virus in the absence of plasma or monoclonal antibodies. The half-maximal neutralization titers for plasma (NT_50_) or half-maximal and 90% inhibitory concentrations for monoclonal antibodies (IC_50_ and IC_90_) were determined using four-parameter nonlinear regression (least squares regression method without weighting; constraints: top=1, bottom=0) (GraphPad Prism).

### Biotinylation of viral protein for use in flow cytometry

Purified and Avi-tagged SARS-CoV-2 RBD or SARS-CoV-2 RBD KEN mutant (K417N, E484K, N501Y) was biotinylated using the Biotin-Protein Ligase-BIRA kit according to manufacturer’s instructions (Avidity) as described before ^3^. Ovalbumin (Sigma, A5503-1G) was biotinylated using the EZ-Link Sulfo-NHS-LC-Biotinylation kit according to the manufacturer’s instructions (Thermo Scientific). Biotinylated ovalbumin was conjugated to streptavidin-BV711 (BD biosciences, 563262) and RBD to streptavidin-PE (BD Biosciences, 554061) and streptavidin-AF647 (Biolegend, 405237) ^3^.

### Flow cytometry and single cell sorting

Single-cell sorting by flow cytometry was described previously ^3^. Briefly, peripheral blood mononuclear cells were enriched for B cells by negative selection using a pan-B-cell isolation kit according to the manufacturer’s instructions (Miltenyi Biotec, 130-101-638). The enriched B cells were incubated in FACS buffer (1× PBS, 2% FCS, 1 mM EDTA) with the following anti-human antibodies (all at 1:200 dilution): anti-CD20-PECy7 (BD Biosciences, 335793), anti-CD3-APC-eFluro 780 (Invitrogen, 47-0037-41), anti-CD8-APC-eFluor 780 (Invitrogen, 47-0086-42), anti-CD16-APC-eFluor 780 (Invitrogen, 47-0168-41), anti-CD14-APC-eFluor 780 (Invitrogen, 47-0149-42), as well as Zombie NIR (BioLegend, 423105) and fluorophore-labelled RBD and ovalbumin (Ova) for 30 min on ice. Single CD3-CD8-CD14-CD16-CD20+Ova-RBD-PE+RBD-AF647+ B cells were sorted into individual wells of 96-well plates containing 4 μl of lysis buffer (0.5× PBS, 10 mM DTT, 3,000 units/ml RNasin Ribonuclease Inhibitors (Promega, N2615) per well using a FACS Aria III and FACSDiva software (Becton Dickinson) for acquisition and FlowJo for analysis. The sorted cells were frozen on dry ice, and then stored at −80 °C or immediately used for subsequent RNA reverse transcription. For B cell phenotype analysis, in addition to above antibodies, B cells were also stained with following anti-human antibodies: anti-IgG-PECF594 (BD biosciences, 562538), anti-IgM-AF700 (Biolegend, 314538), anti-IgA-Viogreen (Miltenyi Biotec, 130-113-481).

### Antibody sequencing, cloning and expression

Antibodies were identified and sequenced as described previously ^3^. In brief, RNA from single cells was reverse-transcribed (SuperScript III Reverse Transcriptase, Invitrogen, 18080-044) and the cDNA stored at −20 °C or used for subsequent amplification of the variable IGH, IGL and IGK genes by nested PCR and Sanger sequencing. Sequence analysis was performed using MacVector. Amplicons from the first PCR reaction were used as templates for sequence- and ligation-independent cloning into antibody expression vectors. Recombinant monoclonal antibodies were produced and purified as previously described ^3^.

### Biolayer interferometry

Biolayer interferometry assays were performed as previously described ^3^. Briefly, we used the Octet Red instrument (ForteBio) at 30 °C with shaking at 1,000 r.p.m. Epitope-binding assays were performed with protein A biosensor (ForteBio 18-5010), following the manufacturer’s protocol ‘classical sandwich assay’. (1) Sensor check: sensors immersed 30 s in buffer alone (kinetics buffer 10x ForteBio 18-1105 diluted 1x in PBS1x). (2) Capture first antibody: sensors immersed 10 min with Ab1 at 30 μg/ml. (3) Baseline: sensors immersed 30 s in buffer alone. (4) Blocking: sensors immersed 5 min with IgG isotype control at 50 μg/ml. (6) Antigen association: sensors immersed 5 min with RBD at 100 μg/ml. (7) Baseline: sensors immersed 30 s in buffer alone. (8) Association Ab2: sensors immersed 5 min with Ab2 at 30 μg/ml. Curve fitting was performed using the Fortebio Octet Data analysis software (ForteBio). Affinity measurement of anti-SARS-CoV-2 IgGs binding were corrected by subtracting the signal obtained from traces performed with IgGs in the absence of WT RBD. The kinetic analysis using protein A biosensor (ForteBio 18-5010) was performed as follows: (1) baseline: 60sec immersion in buffer. (2) loading: 200sec immersion in a solution with IgGs 30 μg/ml. (3) baseline: 200sec immersion in buffer. (4) Association: 300sec immersion in solution with WT RBD at 200, 100, 50 or 25 μg/ml (5) dissociation: 600sec immersion in buffer. Curve fitting was performed using a fast 1:1 binding model and the Data analysis software (ForteBio). Mean *K_D_* values were determined by averaging all binding curves that matched the theoretical fit with an R^2^ value ≥ 0.8.

### Plasma antibody avidity assay

The plasma SARS-CoV-2 antibody avidity assay were performed as previously described ^47^.

### Computational analyses of antibody sequences

Antibody sequences were trimmed based on quality and annotated using Igblastn v.1.14. with IMGT domain delineation system. Annotation was performed systematically using Change-O toolkit v.0.4.540 ^48^. Heavy and light chains derived from the same cell were paired, and clonotypes were assigned based on their V and J genes using in-house R and Perl scripts (Fig. 2d). All scripts and the data used to process antibody sequences are publicly available on GitHub (https://github.com/stratust/igpipeline).

The frequency distributions of human V genes in anti-SARS-CoV-2 antibodies from this study was compared to 131,284,220 IgH and IgL sequences generated by ^49^ and downloaded from cAb-Rep^50^, a database of human shared BCR clonotypes available at https://cab-rep.c2b2.columbia.edu/. Based on the 91 distinct V genes that make up the 6902 analyzed sequences from Ig repertoire of the 10 participants present in this study, we selected the IgH and IgL sequences from the database that are partially coded by the same V genes and counted them according to the constant region. The frequencies shown in (Extended data Fig. 4) are relative to the source and isotype analyzed. We used the two-sided binomial test to check whether the number of sequences belonging to a specific IgHV or IgLV gene in the repertoire is different according to the frequency of the same IgV gene in the database. Adjusted p-values were calculated using the false discovery rate (FDR) correction. Significant differences are denoted with stars.

Nucleotide somatic hypermutation and CDR3 length were determined using in-house R and Perl scripts. For somatic hypermutations, IGHV and IGLV nucleotide sequences were aligned against their closest germlines using Igblastn and the number of differences were considered nucleotide mutations. The average mutations for V genes were calculated by dividing the sum of all nucleotide mutations across all participants by the number of sequences used for the analysis.

Immunoglobulins grouped into the same clonal lineage had their respective IgH and IgL sequences merged and subsequently aligned, using TranslatorX v.1.1 ^51^, with the unmutated ancestral sequence obtained from IMGT/V-QUEST reference directory ^52^. GCTree (https://github.com/matsengrp/gctree) ^53^ was further used to perform the phylogenetic trees construction. Each node represents a unique IgH and IgL combination and the size of each node is proportional to the number of identical sequences. The numbered nodes represent the unobserved ancestral genotypes between the germline sequence and the sequences on the downstream branch.

## Competing interests

The Rockefeller University has filed a provisional patent application in connection with this work on which M.C.N.is an inventor (US patent 63/021,387). The patent has been licensed by Rockefeller University to Bristol Meyers Squib. Z.Z. received seed instruments and sponsored research funding from ET Healthcare.

## Data availability statement

Data are provided in SI Tables 1-8. The raw sequencing data and computer scripts associated with Figure 2 and Extended data figure 5 have been deposited at Github (https://github.com/stratust/igpipeline). This study also uses data from “A Public Database of Memory and Naive B-Cell Receptor Sequences” (https://doi.org/10.5061/dryad.35ks2) and from “High frequency of shared clonotypes in human B cell receptor repertoires” (https://doi.org/10.1038/s41586-019-0934-8).

## Code availability statement

Computer code to process the antibody sequences is available at GitHub (https://github.com/stratust/igpipeline).

## Data presentation

Figures arranged in Adobe Illustrator 2020.

## Acknowledgements

We thank all study participants who devoted time to our research; The Rockefeller University Hospital nursing staff and Clinical Research Support Office and nursing staff. Mayu Okawa Frank, Marissa Bergh, and Robert B. Darnell for SARS-CoV-2 saliva PCR testing. Pamela J. Bjorkman and all members of the M.C.N. laboratory for helpful discussions and Maša Jankovic for laboratory support. This work was supported by NIH grant P01-AI138398-S1 (M.C.N., C.M.R and P.J.B.) and 2U19AI111825 (M.C.N. and C.M.R); George Mason University Fast Grants to C.M.R., 3 R01-AI091707-10S1 to C.M.R.; The G. Harold and Leila Y. Mathers Charitable Foundation to C.M.R.; NIH grant R37-AI64003 to P.D.B.; NIH grant R01AI78788 to T.H.; We thank Dr. Jost Vielmetter and the Protein Expression Center in the Beckman Institute at Caltech for expression assistance. C.O.B. is supported by the HHMI Hanna Gray and Burroughs Wellcome PDEP fellowships. C.G. was supported by the Robert S. Wennett Post-Doctoral Fellowship, in part by the National Center for Advancing Translational Sciences (National Institutes of Health Clinical and Translational Science Award program, grant UL1 TR001866), and by the Shapiro-Silverberg Fund for the Advancement of Translational Research. P.D.B. and M.C.N. are Howard Hughes Medical Institute Investigators. F.M. is supported by the Bulgari Women & Science Fellowship in COVID-19 Research.

## Author Contributions

P.D.B., T.H., C.M.R., and M.C.N. conceived, designed and analyzed the experiments. M. Caskey and C.G. designed clinical protocols. Z.W., F.M., D.S.B., S.F., C.V., H-H.H., C.O.B., A.C., F.S., J.D.S., E.B., L.A., J.Y., M.J. and Z.Z. carried out experiments. A.G. and M. Cipolla produced antibodies. D.S.B., M.D., M.T., K.G.M., C.G. and M. Caskey recruited participants, executed clinical protocols and processed samples. T.Y.O. and V.R. performed bioinformatic analysis. Z.W., F.M., D.S.B., C.G. and M.C.N. wrote the manuscript with input from all co-authors.

## Extended Data Figures

**Extended Data Fig. 1.**
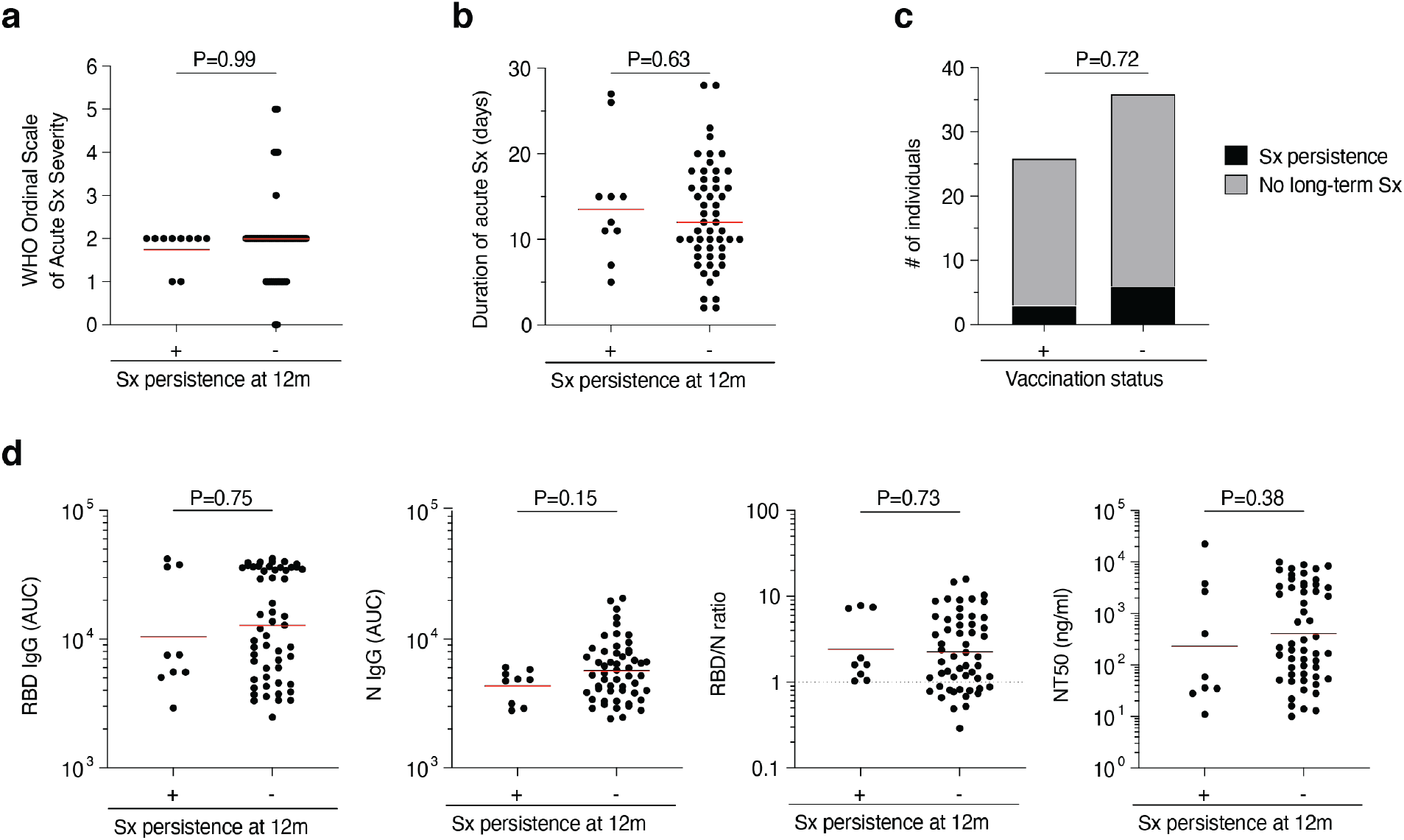
Clinical correlations. **a-d,** Association of persistence of symptoms (Sx) 12 months after infection with various clinical and serological parameters in our cohort of individuals who recovered from COVID-19 (n=63). **a-b,** Acute disease severity as assessed with the WHO Ordinal Scale of Clinical Improvement (**a**, p=0.99) and duration of acute phase symptoms (**b**, p=0.63) in individuals reporting persistent symptoms (+) compared to individuals who are symptom-free (-) 12 months post-infection. **c,** Proportion of individuals reporting persistent symptoms (black area) compared to individuals who are symptom-free (grey area) 12 months after infection grouped by vaccination status (p=0.72). **d,** Anti-RBD IgG (p=0.75), anti-N IgG (p=0.15), the RBD/N IgG ratio (p=0.73), and NT50 titers (p=0.38) at 12 months after infection in individuals reporting persistent symptoms (+) compared to individuals who are symptom-free (-) 12 months post-infection. Statistical significance was determined using the two-tailed Mann-Whitney test in **a, b** and **d**, and using the two-sided Fisher’s exact test in **c**.

**Extended Data Fig. 2:**
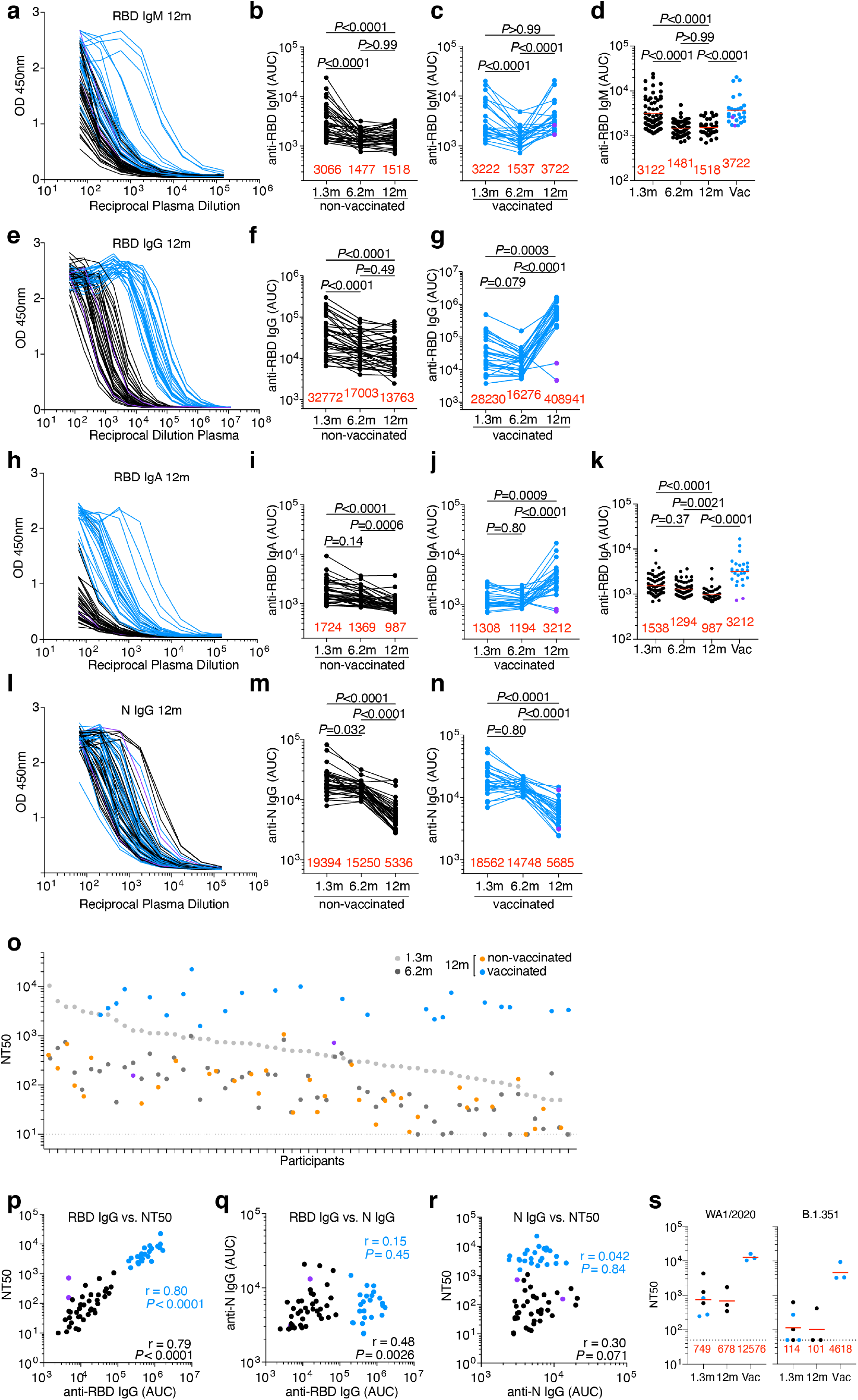
Plasma activity. **a-h,** ELISA results for plasma against SARS-CoV-2 RBD 12 months after infection (n=63). Non-vaccinated individuals are depicted with black circles and lines, and vaccinated individuals are depicted in blue throughout. Two outlier individuals who received their first dose of vaccine 24-48 hours before sample collection is depicted as purple circles. **a-n,** IgM (**a-d**) IgG (**e-g**) and IgA (**h-k**) antibody binding to SARS-CoV-2 RBD and IgG binding to N (**l-n**) 12 months after infection. **a, e, h, I,** ELISA curves from non-vaccinated (black lines) individuals, as well as individuals who received one or two doses (blue lines) of a COVID-19 mRNA vaccine (left panels). Area under the curve (AUC) over time in non-vaccinated (**b, f, i** and **m**) and vaccinated individuals (**c, g, j,** and **n**). Lines connect longitudinal samples. **d, k,** Boxplots showing AUC values of all 63 individuals, as indicated. **o,** ranked average NT50 at 1.3 months (light grey) and 6.2 months (dark grey), as well as at 12 months for non-vaccinated (orange) individuals, and individuals who received one or two doses (blue circles) of a COVID-19 mRNA vaccine, respectively. Two individuals who received their first dose of vaccine 24-48 hours before sample collection is depicted in purple. **p-r,** Correlation of serological parameters in non-vaccinated (black circles and black statistics) and vaccinated (blue circles and blue statistics) individuals. Two individuals who received their first dose of vaccine 24-48 hours before sample collection is depicted as purple circles. Correlation of 12-month titers of anti-RBD IgG and NT50 (**p**), anti-RBD IgG and N IgG (**q**), and anti-N IgG and NT50 (**r**). **s,** Plasma neutralizing activity against authentic virus isolates WA1/2020 and B.1.351, as indicated (n=6). Statistical significance was determined using two-sided Friedman test with subsequent Dunn’s multiple comparisons (**b,c,f,g,i,j,m,n**), or two-sided Kruskal-Wallis test with subsequent Dunn’s multiple comparisons (**d** and **k**) or using the Spearman correlation test for the non-vaccinated and vaccinated subgroups independently (**p-r**) or using two-tailed Mann–Whitney test (**s**). Red numbers indicate the geometric mean NT50 at the indicated timepoint. All experiments were performed at least in duplicate.

**Extended Data Fig. 3:**
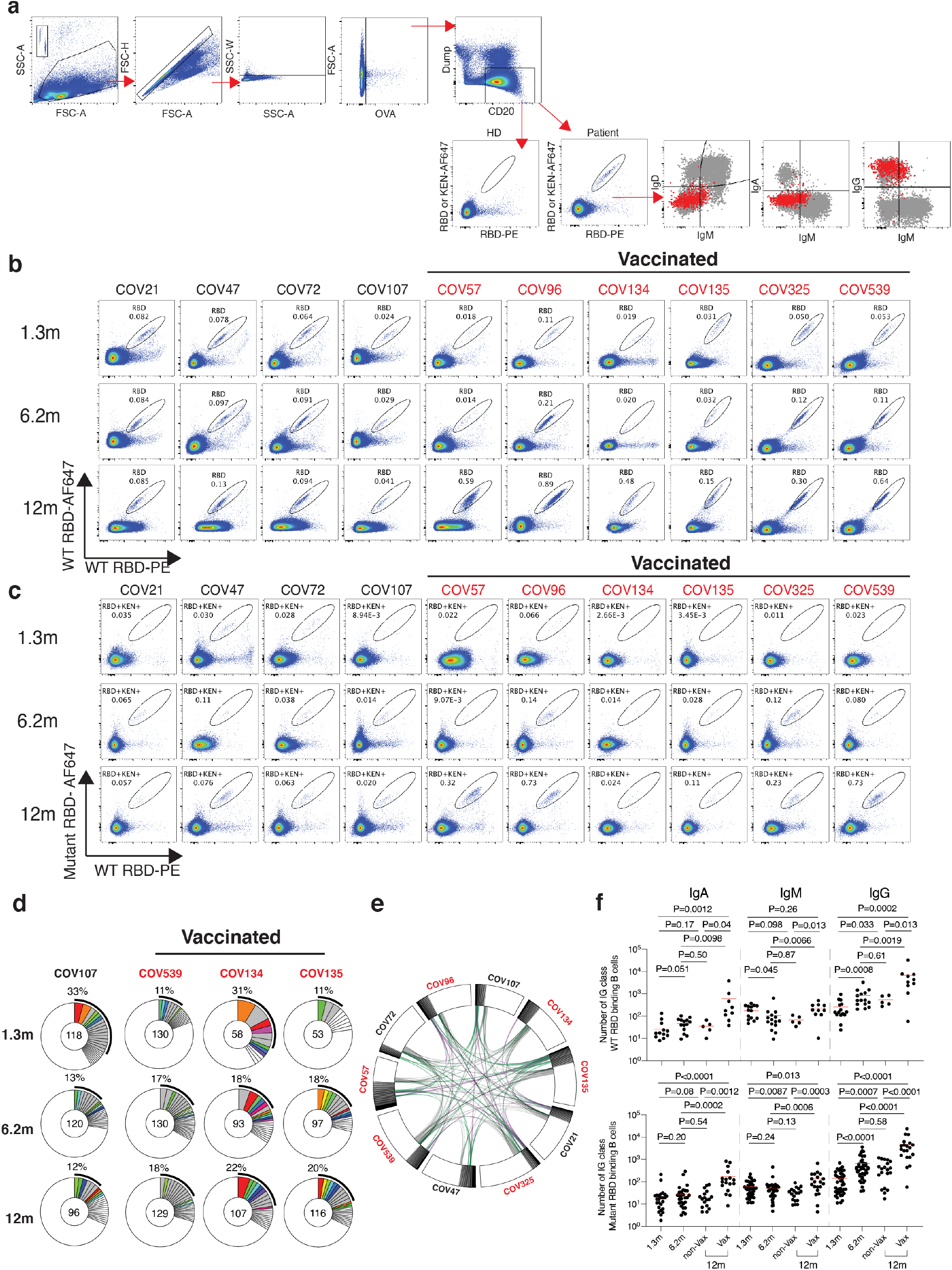
Flow cytometry. **a,** Gating strategy. Gating was on singlets that were CD20^+^ and CD3-CD8-CD16-Ova-. Anti-IgG, IgM, and IgA antibodies were used for B cell phenotype analysis. Sorted cells were RBD-PE^+^ and RBD/KEN-AF647^+^. **b and c,** Flow cytometry showing the percentage of RBD-double positive (**b**) and 647-K417N/E484K/N501Y mutant RBD cross-reactive (**c**) memory B cells from 1.3 or 6- and 12-months post-infection in 10 selected participants. **d**. as in Fig. 2b, Pie charts show the distribution of antibody sequences from 4 individuals after 1.3 ^3^ (upper panel) or 6.2^4^ months (middle panel) or 12 months (lower panel). **e,** Circos plot depicts the relationship between antibodies that share V and J gene segment sequences at both IGH and IGL. Purple, green, and grey lines connect related clones, clones and singles, and singles to each other, respectively. **f,** graph summarizes cell number (indicated in **b** and **c**) (per 2 million B cells) of immunoglobulin class of antigens binding memory B cells in samples obtained at 1.3, 6.2 and 12 months. Each dot is one individual. (Vaccinees, n=20, and non-vaccinees, n=20). Red horizontal bars indicate mean values. Statistical significance was determined using two-sided Kruskal-Wallis test with subsequent Dunn’s multiple comparisons.

**Extended Data Fig. 4:**
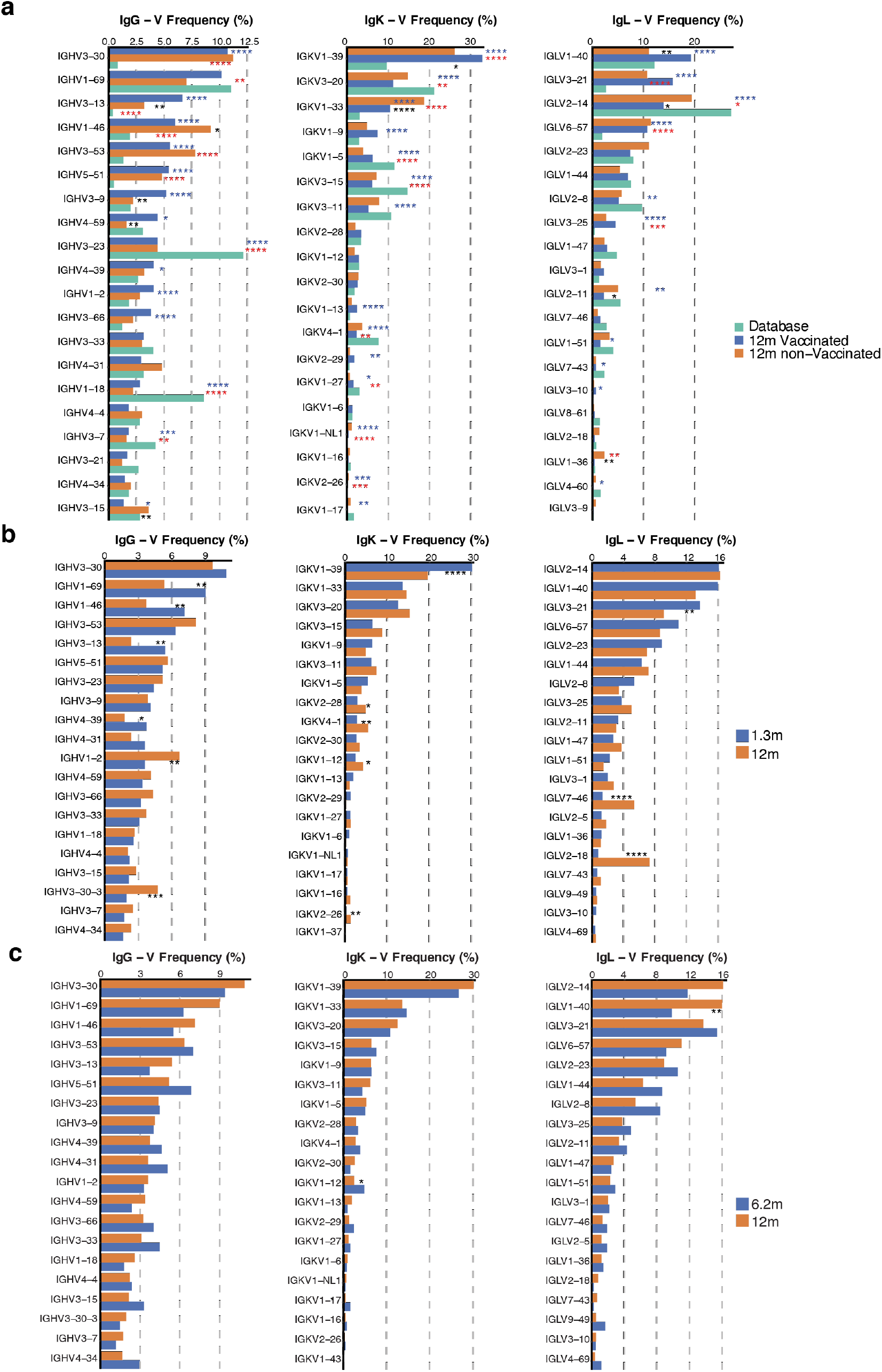
Frequency distribution of human V genes. Graph shows comparison of the frequency distributions of human V genes of anti-SARS-CoV-2 antibodies from donors at 1.3^3^, 6.2^4^, 12 months after infection. **a,** Graph shows relative abundance of human IGVH genes Sequence Read Archive accession SRP010970 (green), convalescent vaccinees (blue), and convalescent non-vaccinees (orange). Statistical significance was determined by two-sided binomial test. **b and c,** same as in **a**, but showing comparison between antibodies from donors at 1.3 months^3^ (**b**), 6.2 month^4^ (**c**), and 12 months after infection. Two-sided binomial tests with unequal variance were used to compare the frequency distributions., significant differences are denoted with stars (* p < 0.05, ** p < 0.01, *** p < 0.001, **** = p < 0.0001).

**Extended Data Fig. 5:**
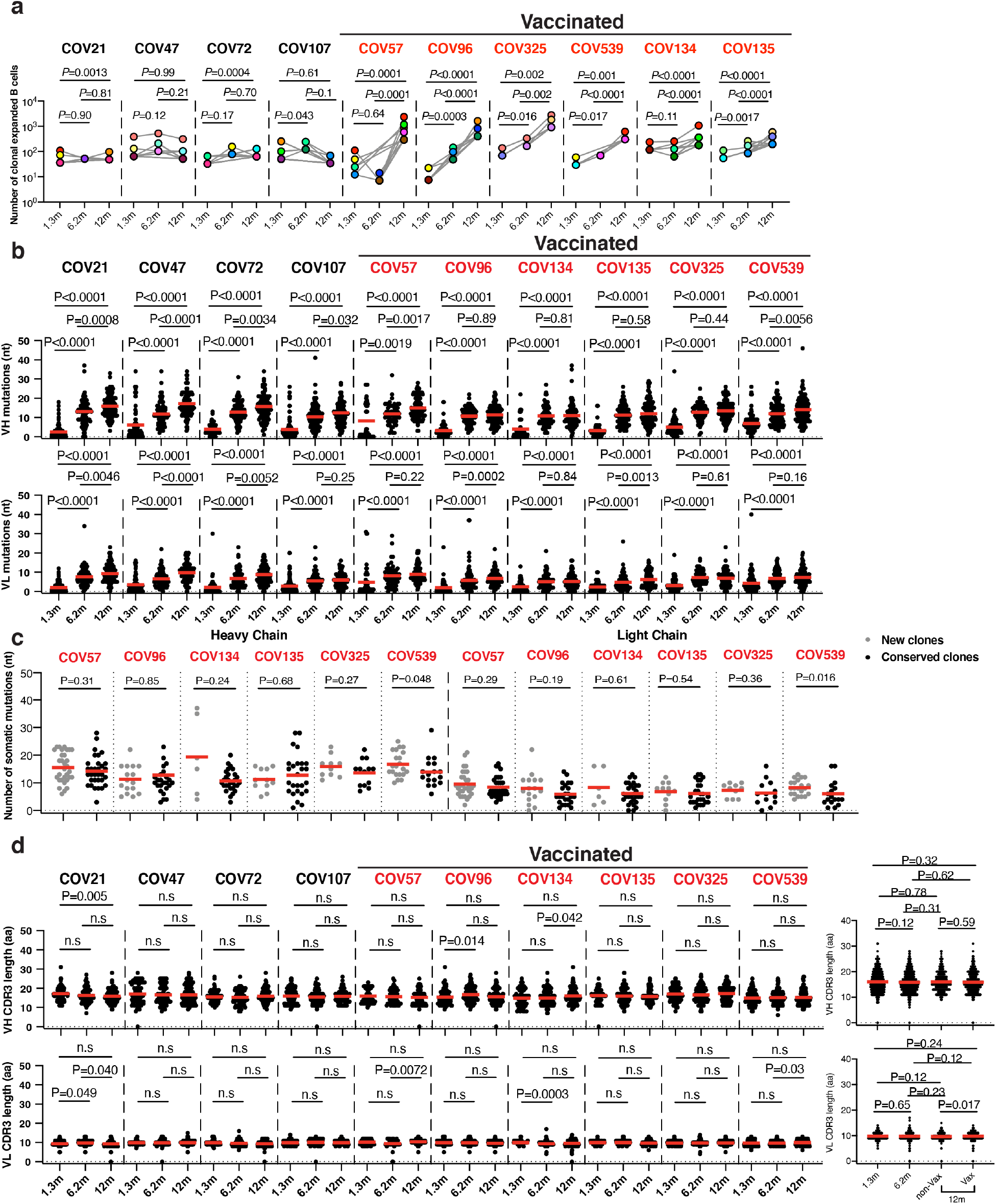
Analysis of anti-RBD antibodies. **a**, Number of clonally expanded B cells (per 10 million B cells) at indicated time points in 10 individuals. Colors indicate shared clones appearing at different time points. Statistical significance was determined using two-tailed Wilcoxon matched-pairs signed rank test. Vaccinees are marked in red. Statistical significance was determined using Wilcoxon matched-pairs signed rank tests. Vaccinees are marked in red. **b.** Number of somatic nucleotide mutations in the IGVH (top) and IGVL (bottom) in antibodies obtained after 1.3 or 6.2 or 12 months from the indicated individual. **c.** same as **b**, but graphs show comparison between new clones and conserved clones in 6 vaccinated convalescent individuals at 12 months after infection. **d.** The amino acid length of the CDR3s at the IGVH and IGVL for each individual. Right panel shows all antibodies combined. (1.3m: n=889; 6.2m: n=975; 12m: n=1105, (non-vax: n=417; vax: n=688)). The horizontal bars indicate the mean. Statistical significance was determined using two-sided Kruskal-Wallis test with subsequent Dunn’s multiple comparisons (**a, b** and **d**), or two-tailed Mann-Whitney U-tests (**c**).

**Extended Data Fig. 6:**
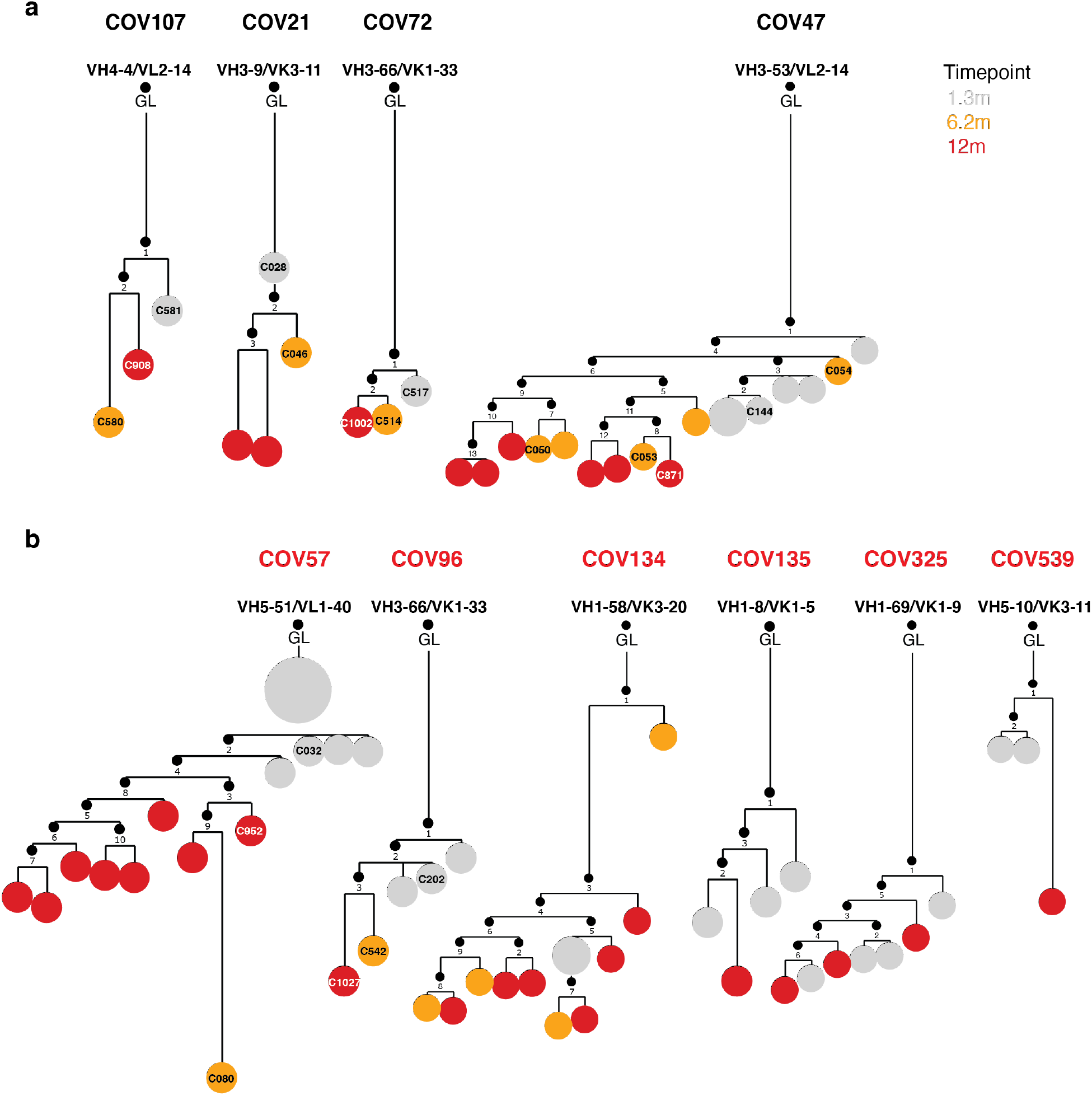
Evolution of anti-SARS-CoV-2 RBD antibody clone. Clonal evolution of RBD-binding memory B cells from ten convalescent individuals, **a**, phylogenetic tree graph shows clones from convalescent non-vaccinees, **b**, same as **a**, but from convalescent vaccinees. Numbers refer to mutations compared to the preceding vertical node. Colors indicate timepoint; grey, orange and red represent 1.3, 6 and 12 months respectively, black dots indicate inferred nodes, and size is proportional to sequence copy number; GL = germline sequence.

**Extended Data Fig. 7:**
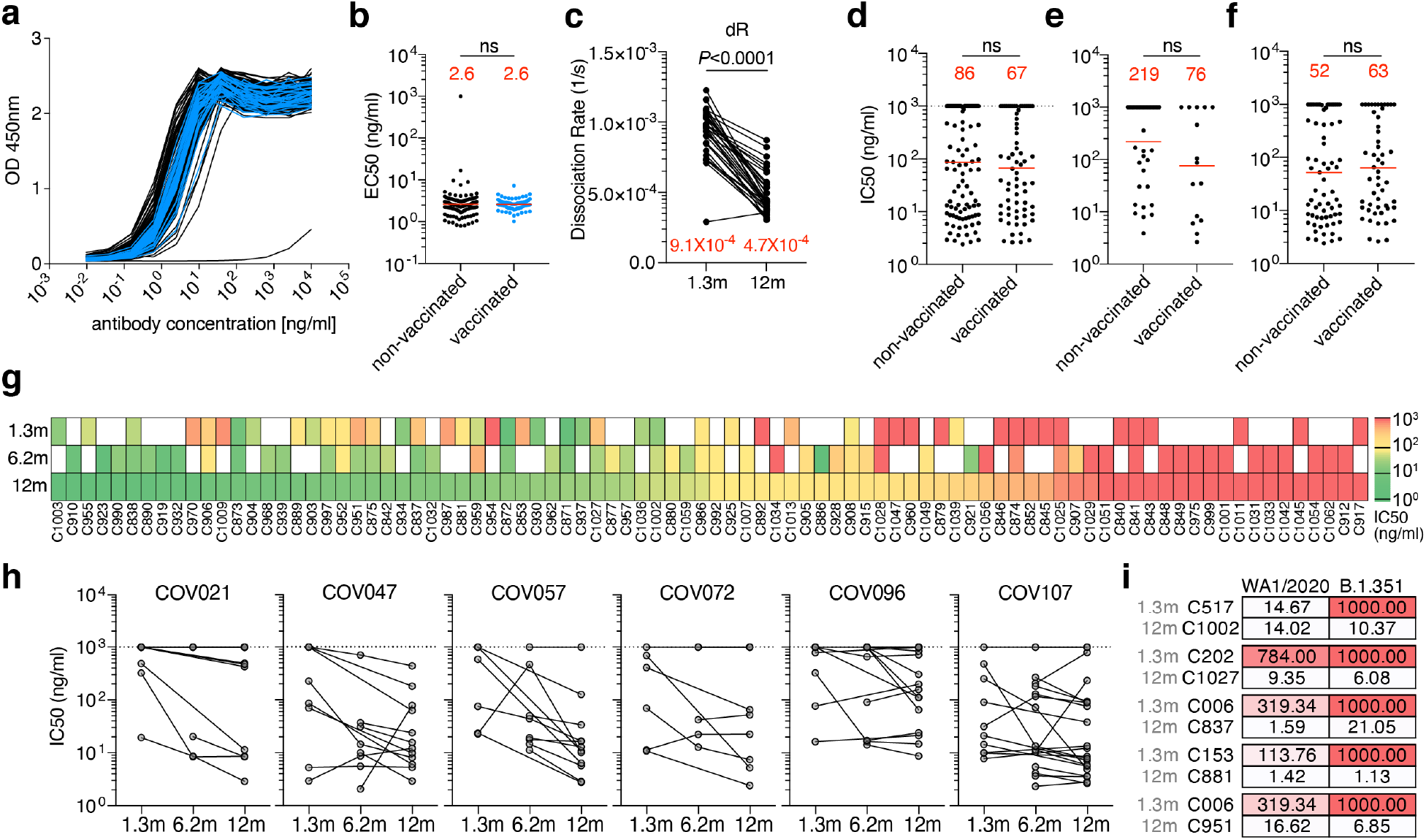
WT RBD binding and pseudovirus neutralization. **a,b,** Binding curves (**a**) and EC_50_ dot plot (**b**) of mAbs isolated from non-vaccinated (black curves and dots) and from vaccinated (blue curves and dots) convalescents individuals 12 months after infection (p=0.74). **c,** Avidity (dissociation rate) measuring plasma reactivity to RBD at the 1.3- and 12-month follow-up visit (n=33). **d-f,** IC_50_ values of mAbs isolated 12 months after infection from non-vaccinated and vaccinated individuals. **d** shows all 12-month antibodies irrespective of clonality, **e** shows singlets only, and **f** shows only antibodies belonging to a clone or shared over time. Statistical significance in **b, d-f** was determined using the two-tailed Mann-Whitney test; **c** was determined using two-tailed Wilcoxon test. The geometric mean EC_50_ and IC_50_ are indicated in red. **g.** Heat map shows the neutralizing activity of clonally related antibodies against wt-SARS-CoV-2 over time. White tiles indicate no clonal relative at the respective time point. Clones are ranked from left to right by the potency of the 12-month progeny antibodies which are denoted below the tiles. **h,** IC_50_ values of shared clones of mAbs cloned from B cells from the initial 1.3- and 6.2, as well as 12-month follow-up visit, divided by participant, as indicated. Lines connect clonal antibodies shared between time points. Antibodies with IC50>1000ng/ml are plotted at 1000 ng/ml in panels d-h. **i,** IC_50_ values of 5 neutralizing antibody pairs against indicated authentic SARS-CoV-2 WA1/2020 and B.1.351 viruses (n=10). Average EC_50_ and IC_50_ values of two independent experiments are shown.

**Extended Data Fig. 8:**
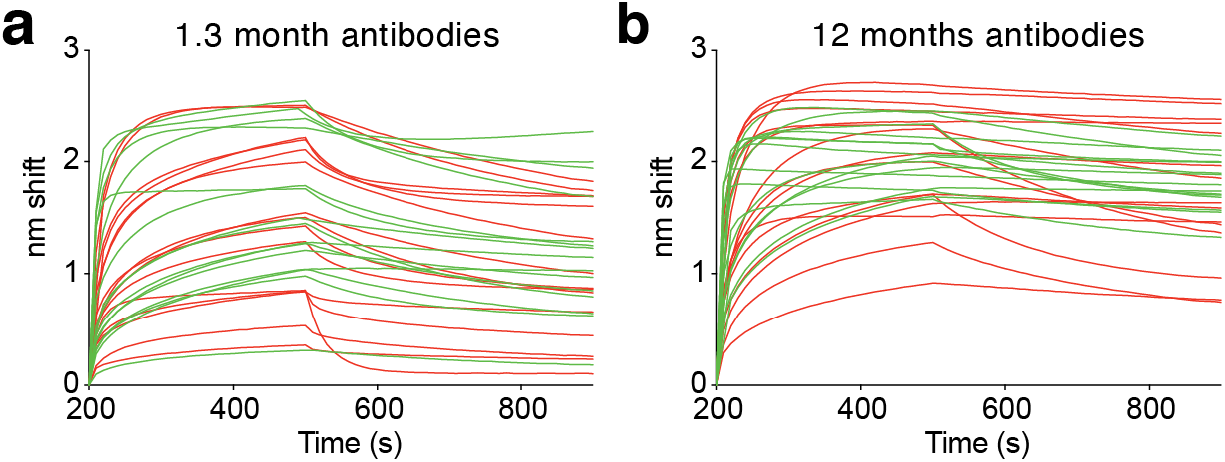
Biolayer interferometry affinity measurements. **a-b,** Graphs depict affinity measurements of neutralizing (green) and non-neutralizing (red) antibodies isolated 1.3 months (**a**) or 12 months (**b**) after infection.

**Extended Data Fig. 9:**
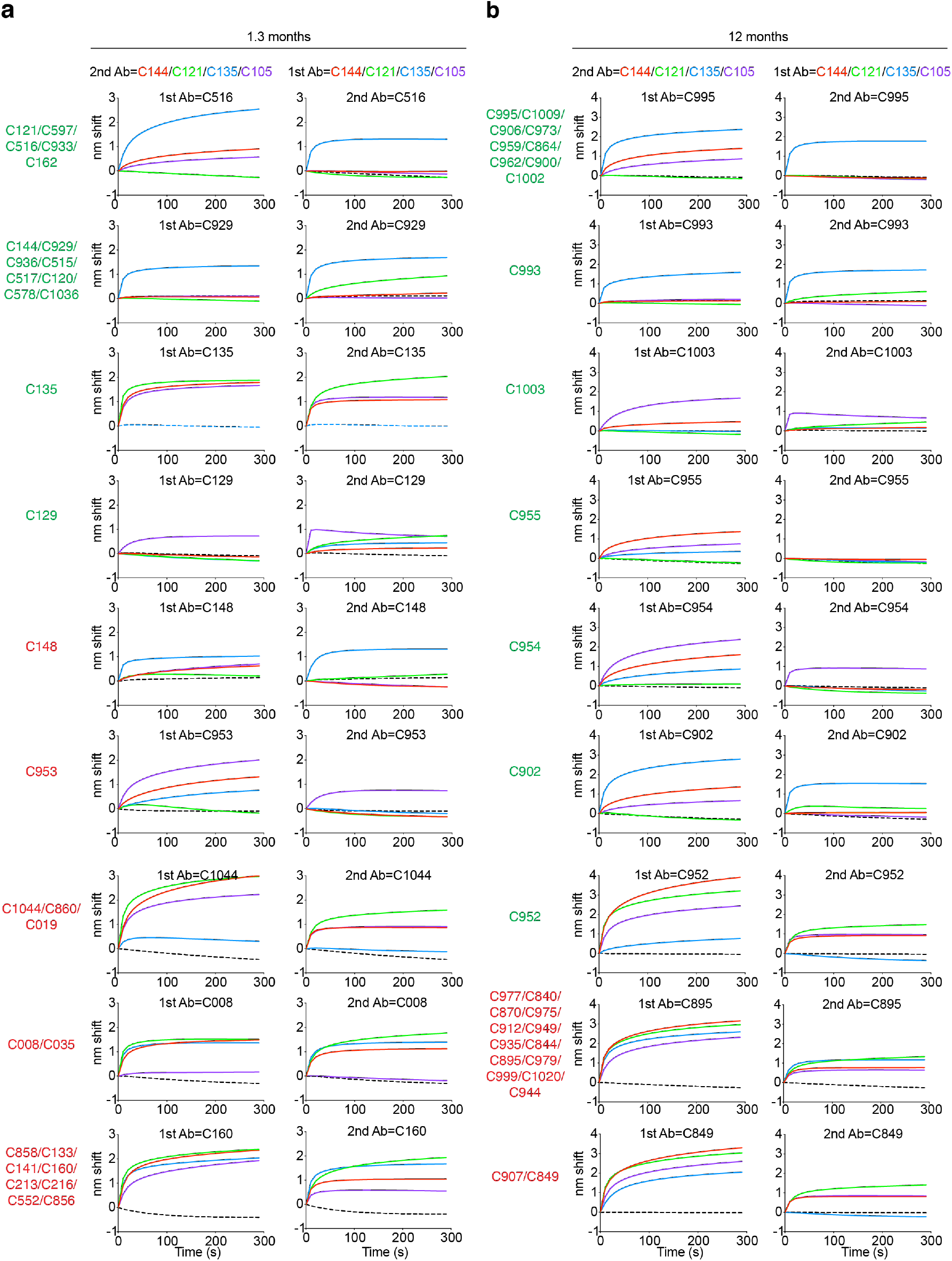
Biolayer interferometry antibody competition experiment. **a-b,** Anti-SARS-CoV-2 RBD antibodies isolated 1.3 (**a**) or 12 months (**b**) after infection were assayed for competition with structurally characterized anti-RBD antibodies by biolayer interferometry experiments as in Fig 4a. Graphs represent the binding of the second antibody (2nd Ab) to preformed first antibody (1st Ab)–RBD complexes. Dotted line denotes when 1st Ab and 2nd Ab are the same. For each antibody group identified in Fig 4c the left graphs represent the binding of the class-representative C144, C121, C135 or C105 ^3,23^(2nd Ab) to the candidate antibody (1st Ab)-RBD complex. The right graphs represent the binding of the candidate antibody (2nd Ab) to the complex of C144-RBD, C121-RBD, C135-RBD or C105-RBD (1st Ab). Antibodies belonging to the same groups are indicated to the left of the respective curves.

## Supplementary Tables

**<u>Supplementary Table 1</u>: Cohort summary**

**<u>Supplementary Table 2</u>: Individual participant characteristics**

**<u>Supplementary Table 3</u>: Antibody sequences from patients is provided as a separate Excel file.**

**<u>Supplementary Table 4</u>: Sequences, half maximal effective concentrations (EC50s) and inhibitory concentrations (IC50s) of the cloned monoclonal antibodies is provided as a separate Excel file.**

**<u>Supplementary Table 5</u>: Binding and Neutralization activity of mAbs against mutant SARS-CoV-2 pseudoviruses.**

**<u>Supplementary Table 6</u>: Neutralization activity of mAbs against mutant SARS-CoV-2 pseudoviruses - Random potently neutralizing antibodies isolated at 1.3 and 12 months**

**<u>Supplementary Table 7</u>: Antibody affinities and neutralization - Clonal pairs isolated at 1.3 and 12 months**

**<u>Supplementary Table 8</u>: Neutralization activity of mAbs against mutant SARS-CoV-2 pseudoviruses - Clonal pairs isolated at 1.3 and 12 months.**

